## Supplementary for "A comprehensive investigation of intracortical and corticothalamic models of the alpha rhythm"

### Supplementary Material

In the following, we provide additional information on various technical details and complementary analyses from our study, which were not included in the main text primarily due to space reasons. These pages cover the derivation of the JR model differential equations (Section S1), the equations of the transfer functions of the four models (Section S2), the derivation of the stability analysis of JR and LW (Section S3), the stability analysis with low noise of JR and LW (Section S4), the phase plane analysis of JR (Section S5), the 4D JR connectivity analyses (Section S6), the comparison of connectivity parameter spaces between JR and MDF (Section S7), the reduced 3D parameter space of MDF (Section S8), the full model equations with a description of their parameters and standard alpha values (Section S9), and finally the additional information on previous models that influenced the four models studied in this paper as well as other existing alpha theories (Section S10). Note that complete code for the generating figures in the following and in the main text is openly available at `github.com/griffithslab/Bastiaens2024_AlphaModels`.

#### S.1 Derivation of the JR Model Equations

The Jansen-Rit and related models are often discussed in terms of a convolution integral for the synaptic impulse response function, as well as the corresponding equivalent second-order differential equation, which is typically what is used in numerical simulations. The mathematical relationship between these is however rarely given in literature sources, and so we provide that here, with a full derivation of the JR differential equation from its impulse response, using the Laplace transform as it simplifies convolution operations by turning them into algebraic manipulations in the Laplace domain.

The synaptic impulse response is defined as an alpha function, which is described by the following equations:

$$h(t) = \begin{cases} \alpha\beta t e^{-\beta t}, & t \geq 0 \\ 0 & \text{otherwise} \end{cases} \tag{1}$$

with $\alpha$ as the maximum amplitude of the PSP and $\beta$ the rate constant parameter. The first step consists of finding the Laplace transform of $h(t)$, denoted as $H(s)$, which is defined as follows:

$$\begin{aligned}
H(s) = \mathcal{L}\{h(t)\} &= \int_0^\infty h(t)e^{-st}\,dt & (2)\\
&= \int_0^\infty \alpha\beta t e^{-\beta t} e^{-st}\,dt & (3)\\
&= \int_0^\infty \alpha\beta t e^{(-\beta-s)t}\,dt & (4)\\
&= \lim_{b\to\infty}\left[\int_0^b \alpha\beta t e^{(-\beta-s)t}\,dt\right] & (5)\\
&= \lim_{b\to\infty}\left(\left[\alpha\beta t \frac{1}{-\beta-s} e^{(-\beta-s)t}\right]_0^b - \int_0^b \frac{\alpha\beta}{-\beta-s} e^{(-\beta-s)t}\,dt\right) & (6)\\
&= \frac{\alpha\beta}{-\beta-s}\lim_{b\to\infty}\left(b e^{(-\beta-s)b} - \int_0^b e^{(-\beta-s)t}\,dt\right) & (7)\\
&= \frac{\alpha\beta}{\beta+s}\lim_{b\to\infty}\int_0^b e^{(-\beta-s)t}\,dt & (8)\\
&= \frac{\alpha\beta}{\beta+s}\lim_{b\to\infty}\left[\frac{1}{-\beta-s} e^{(-\beta-s)t}\right]_0^b & (9)\\
&= \frac{\alpha\beta}{(\beta+s)^2}\lim_{b\to\infty}\left[1 - e^{(-\beta-s)b}\right] & (10)\\
&= \frac{\alpha\beta}{(\beta+s)^2} & (11)
\end{aligned}$$

Now, with an expression for $H(s)$ in the Laplace domain, and given that $y(t)$ is equal to the convolution of $h(t)$ and $x(t)$, we can represent this relationship in the Laplace domain as:

$$\begin{aligned} Y(s) &= X(s)H(s) & (12) \\ Y(s) &= X(s)\frac{\alpha\beta}{(\beta+s)^2} & (13) \\ (\beta+s)^2 Y(s) &= X(s)\alpha\beta & (14) \\ s^2Y(s) + \beta^2 Y(s) + 2\beta s Y(s) &= \alpha\beta X(s) & (15) \\ s^2 Y(s) &= \alpha\beta X(s) - 2\beta s Y(s) - \beta^2 Y(s) & (16) \end{aligned}$$

Since $s^2Y(s)$ corresponds to the second derivative in the time domain, translating equation (40) back into the time domain, we obtain:

$$\ddot{y}(t) = \alpha\beta x(t) - 2\beta\dot{y}(t) - \beta^2 y(t) \tag{17}$$

This corresponds to the commonly used JR second-order differential equation, which can be rewritten in the form of two first-order ODE's:

$$\begin{aligned} \dot{y}(t) &= z(t) & (18) \\ \dot{z}(t) &= \alpha\beta x(t) - 2\beta z(t) - \beta^2 y(t) & (19) \end{aligned}$$

with $y(t)$ representing the average postsynaptic membrane potential (output of the PSP block).

#### S.2 Transfer Function of the Models

In addition to comparing the numerical simulation outputs, we also simulated the analytical (or linearized) power spectra of the models. These analytical models are used for stability analysis, offering clear insights into how a system responds to changes in inputs. Moreover, they provide explicit solutions, enhancing our understanding of the system's behavior. The transfer functions for both the JR and MDF models are derived from graph control analysis. The equations for the LW model are sourced from Hartoyo et al. (2019), while those for the RRW model are from Robinson et al. (2002).

JR Transfer Function:

$$T(s) = \frac{G_e}{1 + K_s^2 G_e (C_3 C_4 G_i - C_1 C_2 G_e)}$$

with:

$$G_e = \frac{Aa^{-1}}{(a^{-1}+s)^2},\ G_i = \frac{Bb^{-1}}{(a^{-1}+s)^2},\ \text{and } K_s = \frac{e_0 r}{2}$$

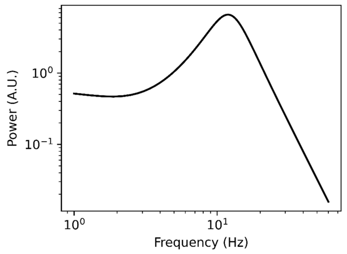


MDF Transfer Function:

$$T(s) = \frac{G_e^2 K_s \gamma_2\,(1 + K_s G_i \gamma_5)}{1 + K_s G_i \gamma_5 + G_e G_i K_s^2 \gamma_3 \gamma_4 - G_e^2 K_s^2 \gamma_2 \gamma_1\,(1 + K_s G_i \gamma_5)}$$

with:

$$G_e = \frac{H_e \kappa_e^{-1}}{(\kappa_e^{-1}+s)^2},\ G_i = \frac{H_i \kappa_i^{-1}}{(\kappa_i^{-1}+s)^2},\ \text{and } K_s = \frac{e_0 r}{2}$$

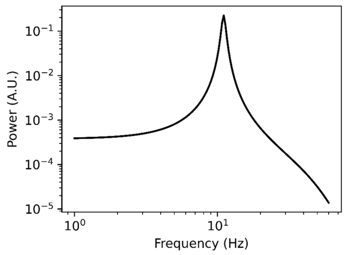


LW Transfer Function:

$$T(s) = \frac{H_1}{1 + H_1 H_2}$$

with:

$$H_1 = \frac{Q_{ee}}{Q_{ee} A_{ee} - K_{ei} Y_e},\ H_2 = \frac{Q_{ie} Q_{ei} A_{ei} A_{ie}}{Q_{ee}(K_{ie} Y_i - Q_{ii} A_{ii})},\ Q_{xy} = -\frac{\psi_{xy}(V_y)}{\tau_y},$$

$$A_{xy} = -\Gamma_x \gamma_x e N_{xy}^b S_x'(V_x),\qquad I_{xy} = \frac{\Gamma_x e}{\gamma_x} N_{xy}^b S_x(V_x) + \frac{\Gamma_x e}{\gamma_x} p_{xy},$$

$$Y_x = (s + \gamma_x)^2,\qquad K_{xy} = s + \frac{1}{\tau_x}\left(1 + \frac{I_{xx}}{|V_x^{eq} - V_x^{rest}|} + \frac{I_{yx}}{|V_y^{eq} - V_x^{rest}|}\right)$$

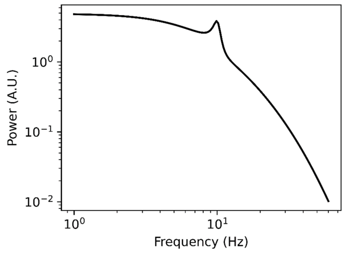


RRW Transfer Function:

$$T(S) = \frac{G_{es} G_{sn} L^2 e^{s t_0/2}}{(1 - G_{ei} L)(1 - G_{srs} L^2)(q^2 r_e^2)}$$

with:

$$L = \left(1 - \frac{s}{\alpha}\right)^{-1} \left(1 - \frac{s}{\beta}\right)^{-1}$$

$$q^2 r_e^2 = \left(1 - \frac{s}{\gamma_e}\right)^2 - \frac{L}{1 - G_{ei} L}\left(G_{ee} + \frac{G_{ese} L + G_{esre} L^2}{1 - G_{srs} L^2} e^{s t_0}\right)$$

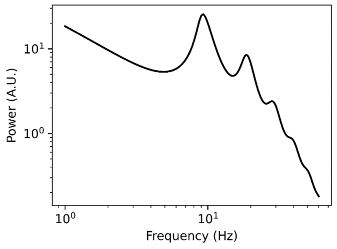


**Figure S1.** ***Transfer function of each of the four studied models (JR, MDF, LW, and RRW) with the corresponding analytical power spectra*** The transfer function of JR and MDF were derived using graph control analysis. The derivations for LW transfer function are sourced from Hartoyo et al. (2019), and for RRW from Robinson et al. (2002)

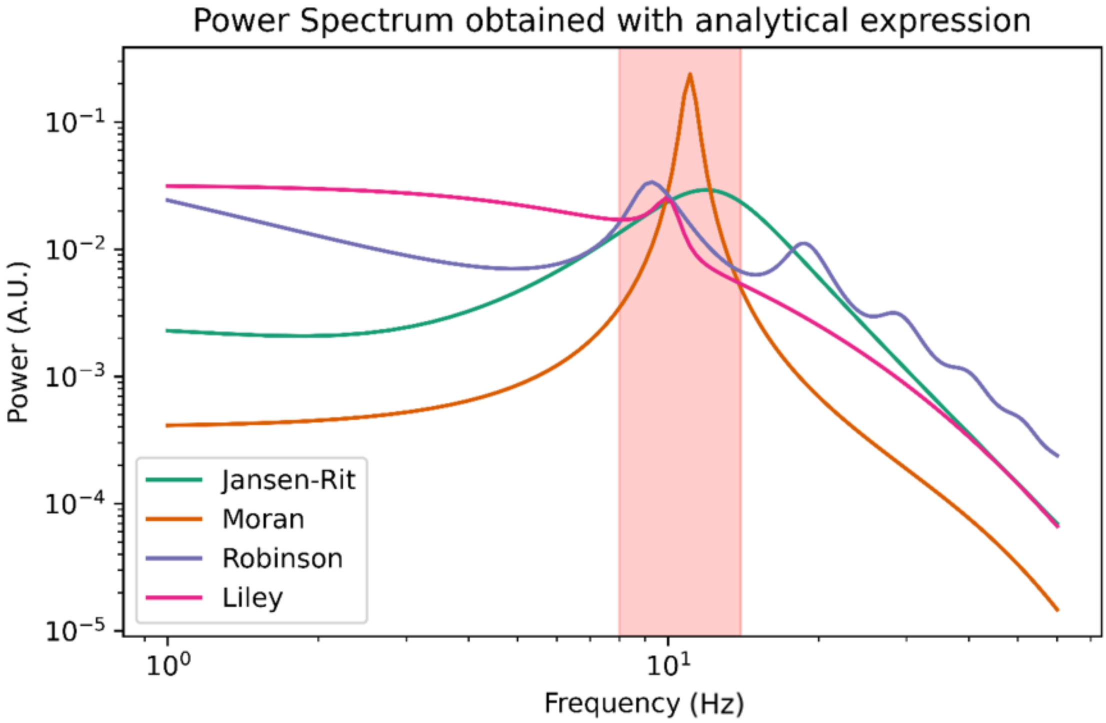


**Figure S2.** ***Analytical power spectra of the four models together.*** After linearization, the models continue to generate alpha rhythms with a peak between 8-12 Hz (represented by the red zone). Among the models, RRW is unique in capturing the harmonics at higher frequencies in its linearized form. JR exhibits a broader peak compared to the others, while LW has a lower peak height.

#### S.3 Derivation of Stability Analysis for JR and LW

For JR, similar to Grimbert and Faugeras (2006), the fixed points are determined by setting the derivatives to 0. With some manipulations, the equilibrium points in the $(C, y_1 - y_2)$ plane with $y = y_1 - y_2$ are equal to:

$$y = \frac{A}{a}p + \frac{A}{a}C_2 S(\frac{A}{a}C_1 S(y) - \frac{B}{b}C_4 S(\frac{A}{a}C_3 S(y)) \tag{20}$$

The stability of the fixed points is then defined using the Jacobian matrix

$$\mathbf{Y}_{i,j} = \begin{bmatrix} \frac{\partial y_0}{\partial y_0} & \frac{\partial y_0}{\partial y_1} & \frac{\partial y_0}{\partial y_2} & \frac{\partial y_0}{\partial y_3} & \frac{\partial y_0}{\partial y_4} & \frac{\partial y_0}{\partial y_5} \\ \frac{\partial y_1}{\partial y_0} & \frac{\partial y_1}{\partial y_1} & \frac{\partial y_1}{\partial y_2} & \frac{\partial y_1}{\partial y_3} & \frac{\partial y_1}{\partial y_4} & \frac{\partial y_1}{\partial y_5} \\ \frac{\partial y_2}{\partial y_0} & \frac{\partial y_2}{\partial y_1} & \frac{\partial y_2}{\partial y_2} & \frac{\partial y_2}{\partial y_3} & \frac{\partial y_2}{\partial y_4} & \frac{\partial y_2}{\partial y_5} \\ \frac{\partial y_3}{\partial y_0} & \frac{\partial y_3}{\partial y_1} & \frac{\partial y_3}{\partial y_2} & \frac{\partial y_3}{\partial y_3} & \frac{\partial y_3}{\partial y_4} & \frac{\partial y_3}{\partial y_5} \\ \frac{\partial y_4}{\partial y_0} & \frac{\partial y_4}{\partial y_1} & \frac{\partial y_4}{\partial y_2} & \frac{\partial y_4}{\partial y_3} & \frac{\partial y_4}{\partial y_4} & \frac{\partial y_4}{\partial y_5} \\ \frac{\partial y_5}{\partial y_0} & \frac{\partial y_5}{\partial y_1} & \frac{\partial y_5}{\partial y_2} & \frac{\partial y_5}{\partial y_3} & \frac{\partial y_5}{\partial y_4} & \frac{\partial y_5}{\partial y_5} \end{bmatrix} = \begin{bmatrix} 0 & 0 & 0 & 1 & 0 & 0 \\ 0 & 0 & 0 & 0 & 1 & 0 \\ 0 & 0 & 0 & 0 & 0 & 1 \\ -a^2 & AaS'(y) & -AaS'(y) & -2a & 0 & 0 \\ AaC_2C_1S'(C_1y_0(y)) & -a^2 & 0 & 0 & -2a & 0 \\ BbC_4C_3S'(C_3y_0(y)) & 0 & -b^2 & 0 & 0 & -2b \end{bmatrix}$$

with $y$ corresponding to the fixed point of interest and $y_0(y) = \frac{A}{a}S(y)$. Stability is then defined by calculating the eigenvalues of the matrix $\mathbf{Y}$ for each fixed point, and looking at the sign of the real part of the eigenvalues. The system is stable if all the eigenvalues have a negative real part. If at least one of the eigenvalues has a positive real part, it is considered as an unstable fixed point.

Using a similar method (estimation of the fixed point, following an assessment of the stability of the fixed points by looking at the real part of the eigenvalues of the Jacobian matrix), the LW equilibrium points' stability was also determined. The full calculation and equations are detailed in the appendix of Hartoyo et al. (2019) and also in Supplementary S.6. Briefly:

The equilibrium point equations can be reduced to:

$$0 = -V_e + V_{er} + \psi_{ee}(V_e)I_{ee} + \psi_i e(V_e)I_{ie} \tag{21}$$

$$0 = -V_i + V_{ir} + \psi_{ei}(V_i)I_{ei} + \psi_{ii}(V_i)I_{ii} \tag{22}$$

with

$$I_{ee} = \frac{\Gamma_e e}{\gamma_e}N_{ee}^{\beta}S(V_e) + \frac{\Gamma_e e}{\gamma_e}p_{ee} \tag{23}$$

$$I_{ei} = \frac{\Gamma_e e}{\gamma_e}N_{ei}^{\beta}S(V_e) + \frac{\Gamma_e e}{\gamma_e}p_{ei} \tag{24}$$

$$I_{ie} = \frac{\Gamma_i e}{\gamma_i}N_{ie}^{\beta}S(V_i) + \frac{\Gamma_i e}{\gamma_i} \tag{25}$$

$$I_{ii} = \frac{\Gamma_i e}{\gamma_i}N_{ii}^{\beta}S(V_i) + \frac{\Gamma_i e}{\gamma_i} \tag{26}$$

The fixed points for $V_e$ and $V_i$ are then estimated by finding the values for which values these two equations intersect.

The Jacobian matrix is:

$$
\mathbf{F}_{i,j} = \begin{bmatrix}
\frac{\partial V_e}{\partial V_e} & \frac{\partial V_e}{\partial V_i} & \frac{\partial V_e}{\partial I_{ee}} & \frac{\partial V_e}{\partial I_{ei}} & \frac{\partial V_e}{\partial I_{ie}} & \frac{\partial V_e}{\partial I_{ii}} & \frac{\partial V_e}{\partial U_{ee}} & \frac{\partial V_e}{\partial U_{ei}} & \frac{\partial V_e}{\partial U_{ie}} & \frac{\partial V_e}{\partial U_{ii}} \\
\frac{\partial V_i}{\partial V_e} & \frac{\partial V_i}{\partial V_i} & \frac{\partial V_i}{\partial I_{ee}} & \frac{\partial V_i}{\partial I_{ei}} & \frac{\partial V_i}{\partial I_{ie}} & \frac{\partial V_i}{\partial I_{ii}} & \frac{\partial V_i}{\partial U_{ee}} & \frac{\partial V_i}{\partial U_{ei}} & \frac{\partial V_i}{\partial U_{ie}} & \frac{\partial V_i}{\partial U_{ii}} \\
\frac{\partial I_{ee}}{\partial V_e} & \frac{\partial I_{ee}}{\partial V_i} & \frac{\partial I_{ee}}{\partial I_{ee}} & \frac{\partial I_{ee}}{\partial I_{ei}} & \frac{\partial I_{ee}}{\partial I_{ie}} & \frac{\partial I_{ee}}{\partial I_{ii}} & \frac{\partial I_{ee}}{\partial U_{ee}} & \frac{\partial I_{ee}}{\partial U_{ei}} & \frac{\partial I_{ee}}{\partial U_{ie}} & \frac{\partial I_{ee}}{\partial U_{ii}} \\
\frac{\partial I_{ei}}{\partial V_e} & \frac{\partial I_{ei}}{\partial V_i} & \frac{\partial I_{ei}}{\partial I_{ee}} & \frac{\partial I_{ei}}{\partial I_{ei}} & \frac{\partial I_{ei}}{\partial I_{ie}} & \frac{\partial I_{ei}}{\partial I_{ii}} & \frac{\partial I_{ei}}{\partial U_{ee}} & \frac{\partial I_{ei}}{\partial U_{ei}} & \frac{\partial I_{ei}}{\partial U_{ie}} & \frac{\partial I_{ei}}{\partial U_{ii}} \\
\frac{\partial I_{ie}}{\partial V_e} & \frac{\partial I_{ie}}{\partial V_i} & \frac{\partial I_{ie}}{\partial I_{ee}} & \frac{\partial I_{ie}}{\partial I_{ei}} & \frac{\partial I_{ie}}{\partial I_{ie}} & \frac{\partial I_{ie}}{\partial I_{ii}} & \frac{\partial I_{ie}}{\partial U_{ee}} & \frac{\partial I_{ie}}{\partial U_{ei}} & \frac{\partial I_{ie}}{\partial U_{ie}} & \frac{\partial I_{ie}}{\partial U_{ii}} \\
\frac{\partial I_{ii}}{\partial V_e} & \frac{\partial I_{ii}}{\partial V_i} & \frac{\partial I_{ii}}{\partial I_{ee}} & \frac{\partial I_{ii}}{\partial I_{ei}} & \frac{\partial I_{ii}}{\partial I_{ie}} & \frac{\partial I_{ii}}{\partial I_{ii}} & \frac{\partial I_{ii}}{\partial U_{ee}} & \frac{\partial I_{ii}}{\partial U_{ei}} & \frac{\partial I_{ii}}{\partial U_{ie}} & \frac{\partial I_{ii}}{\partial U_{ii}} \\
\frac{\partial U_{ee}}{\partial V_e} & \frac{\partial U_{ee}}{\partial V_i} & \frac{\partial U_{ee}}{\partial I_{ee}} & \frac{\partial U_{ee}}{\partial I_{ei}} & \frac{\partial U_{ee}}{\partial I_{ie}} & \frac{\partial U_{ee}}{\partial I_{ii}} & \frac{\partial U_{ee}}{\partial U_{ee}} & \frac{\partial U_{ee}}{\partial U_{ei}} & \frac{\partial U_{ee}}{\partial U_{ie}} & \frac{\partial U_{ee}}{\partial U_{ii}} \\
\frac{\partial U_{ei}}{\partial V_e} & \frac{\partial U_{ei}}{\partial V_i} & \frac{\partial U_{ei}}{\partial I_{ee}} & \frac{\partial U_{ei}}{\partial I_{ei}} & \frac{\partial U_{ei}}{\partial I_{ie}} & \frac{\partial U_{ei}}{\partial I_{ii}} & \frac{\partial U_{ei}}{\partial U_{ee}} & \frac{\partial U_{ei}}{\partial U_{ei}} & \frac{\partial U_{ei}}{\partial U_{ie}} & \frac{\partial U_{ei}}{\partial U_{ii}} \\
\frac{\partial U_{ie}}{\partial V_e} & \frac{\partial U_{ie}}{\partial V_i} & \frac{\partial U_{ie}}{\partial I_{ee}} & \frac{\partial U_{ie}}{\partial I_{ei}} & \frac{\partial U_{ie}}{\partial I_{ie}} & \frac{\partial U_{ie}}{\partial I_{ii}} & \frac{\partial U_{ie}}{\partial U_{ee}} & \frac{\partial U_{ie}}{\partial U_{ei}} & \frac{\partial U_{ie}}{\partial U_{ie}} & \frac{\partial U_{ie}}{\partial U_{ii}} \\
\frac{\partial U_{ii}}{\partial V_e} & \frac{\partial U_{ii}}{\partial V_i} & \frac{\partial U_{ii}}{\partial I_{ee}} & \frac{\partial U_{ii}}{\partial I_{ei}} & \frac{\partial U_{ii}}{\partial I_{ie}} & \frac{\partial U_{ii}}{\partial I_{ii}} & \frac{\partial U_{ii}}{\partial U_{ee}} & \frac{\partial U_{ii}}{\partial U_{ei}} & \frac{\partial U_{ii}}{\partial U_{ie}} & \frac{\partial U_{ii}}{\partial U_{ii}}
\end{bmatrix}
$$

which evaluates to

$$
\mathbf{F}_{i,j} = \begin{bmatrix}
G(V_e) & 0 & \frac{\psi_{ee}(V_e)}{\tau_e} & 0 & \frac{\psi_{ie}(V_e)}{\tau_e} & 0 & 0 & 0 & 0 & 0 \\
G_(V_i) & 0 & \frac{\psi_{ei}(V_i)}{\tau_i} & 0 & \frac{\psi_{ii}(V_i)}{\tau_i} & 0 & 0 & 0 & 0 & 0 \\
0 & 0 & 0 & 0 & 0 & 0 & 1 & 0 & 0 & 0 \\
0 & 0 & 0 & 0 & 0 & 0 & 0 & 1 & 0 & 0 \\
0 & 0 & 0 & 0 & 0 & 0 & 0 & 0 & 1 & 0 \\
0 & 0 & 0 & 0 & 0 & 0 & 0 & 0 & 0 & 1 \\
\Gamma_e\gamma_e e N_{ee}^{\beta} S'(V_e) & 0 & -\gamma_e^2 & 0 & 0 & 0 & -2\gamma_e & 0 & 0 & 0 \\
\Gamma_e\gamma_e e N_{ei}^{\beta} S'(V_e) & 0 & 0 & -\gamma_e^2 & 0 & 0 & 0 & -2\gamma_e & 0 & 0 \\
0 & \Gamma_i\gamma_i e N_{ie}^{\beta} S'(V_i) & 0 & 0 & -\gamma_i^2 & 0 & 0 & 0 & -2\gamma_i & 0 \\
0 & \Gamma_i\gamma_i e N_{ii}^{\beta} S'(V_i) & 0 & 0 & 0 & -\gamma_i^2 & 0 & 0 & 0 & -2\gamma_i
\end{bmatrix}
$$

with

$$
G(V_e) = \frac{1}{\tau_e}\left(-1 - \frac{I_{ee}}{|V_e^e q - V_{er}|} - \frac{I_{ie}}{|V_i^e q - V_{ir}|}\right) \tag{27}
$$

$$
G(V_i) = \frac{1}{\tau_i}\left(-1 - \frac{I_{ei}}{|V_e^e q - V_{ir}|} - \frac{I_{ii}}{|V_i^e q - V_{ir}|}\right) \tag{28}
$$

We then replace $V_e$ and $V_i$ with the equilibrium points computed previously, and the real parts of the eigenvalues of this Jacobian matrix are then examined to assess their stability.

#### S.4 Stability Analysis Low Noise JR and LW of Connectivity E-I parameters

In the main text, we presented the stability analysis of the E-I connectivity parameters with standard input. Our aim was to demonstrate the changes in this analysis when low noise input is introduced, highlighting the differences between the JR and LW models. Specifically, we showed that in the JR model, a Hopf bifurcation occurs at the alpha rhythm. In contrast, the alpha rhythm generated with LW model is noise-driven, as without noise, it behaves as a damped oscillator and reaches a fixed point instead of oscillating.

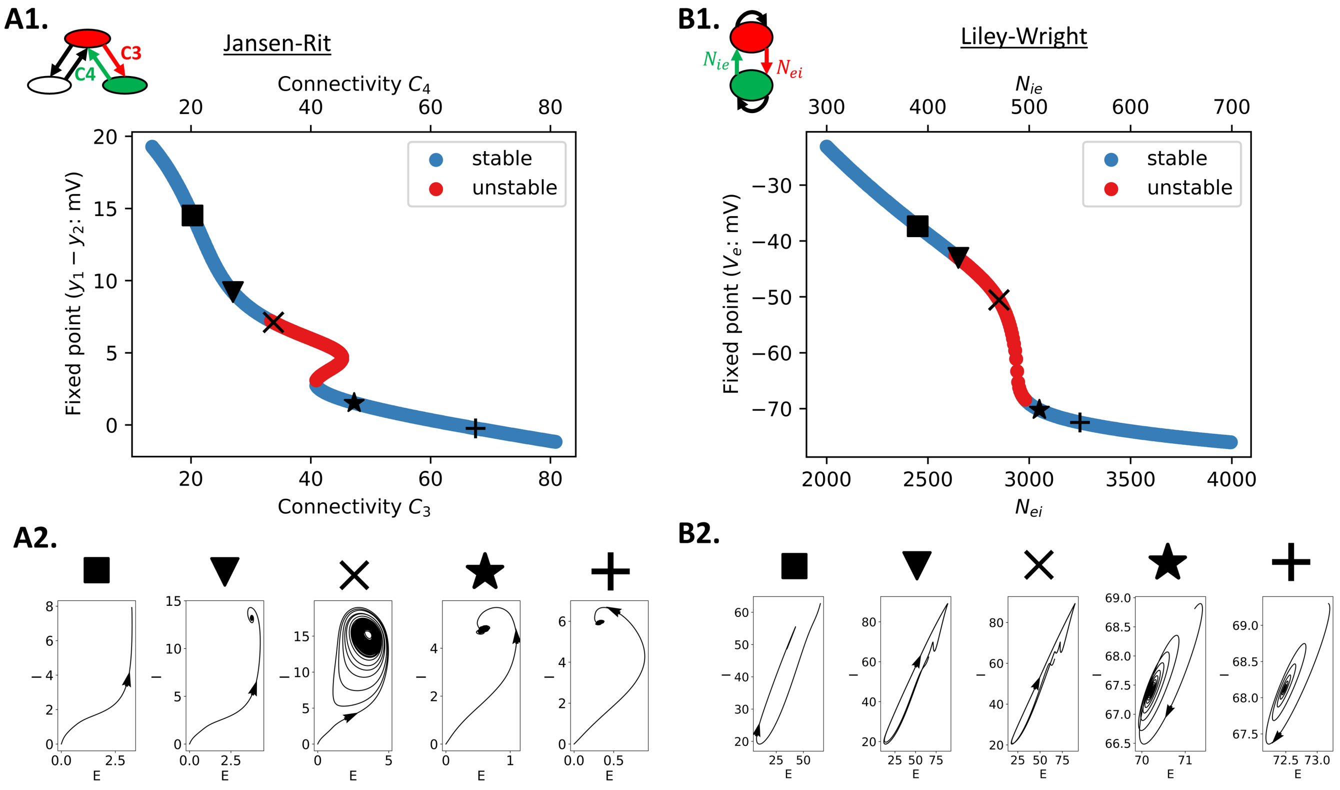


**Figure S3.** ***Fixed points and corresponding phase planes of JR and LW at specific connectivity values with low noise input*** For the JR model, **A1** and **A2** correspond to the fixed points of JR with low noise, and the corresponding phase planes for specific connectivity values. Similarly to JR, the fixed points of LW are presented with low noise and the corresponding phase plane for specific connetivity values in **B1** and **B2**. Unstable fixed points are red, whereas stable fixed points are blue. The cross phase plan in JR correspond to the standard alpha rhythm parameters, whereas the star phase plane in LW correspond to the standard alpha parameters.

#### S.5 Phase plane of JR in 3D

For the stability analyses in Fig. 11, we have only presented the phase plane with the pyramidal and inhibitory population output voltages. Considering the trajectory of the third excitatory neural population activity alongside these can provide a better understanding of the full picture however, as can be seen in Fig. S4.

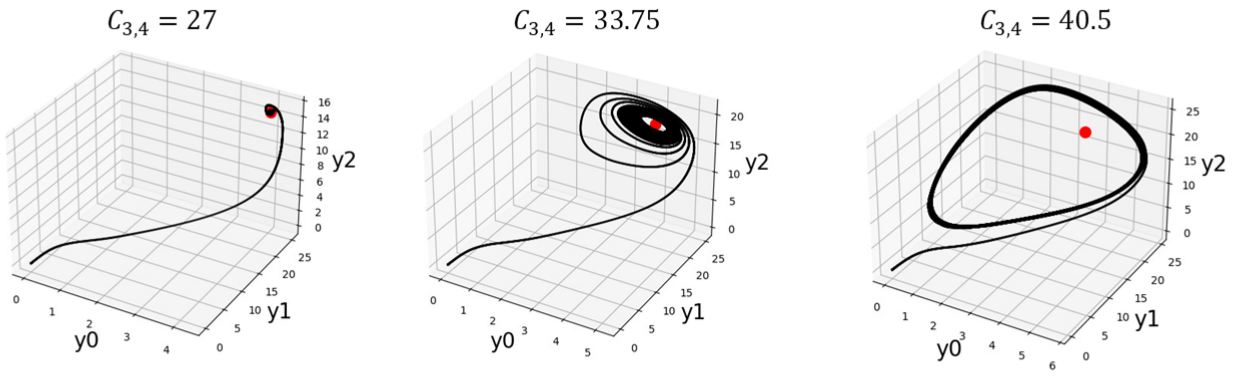


**Figure S4.** ***Phase plane of JR for different E-I connectivity parameters.*** The trajectory of the three neural populations (with $y_0$, $y_1$, and $y_2$ corresponding to the output of the PSP block for the pyramidal cells, excitatory interneurons, and inhibitory interneurons, respectively) can be inferred by examining the stability of their respective fixed point (red). When $C_{3,4} = 27$, the fixed point is stable and no oscillations occur. For $C_{3,4} = 33.5$, the system enters a limit cycle with the oscillation frequency of alpha. Finally, when $C_{3,4} = 40.5$, the limit cycle widens and the frequency of oscillation is reduced.

As seen in our previous phase plane analysis, for specific connectivity parameters, the system either reaches a fixed point or enters a limit cycle defining the frequency of oscillation. The results closely resemble those in Fig. 11, implying that the dynamics primarily involve interactions between the pyramidal and inhibitory populations, with minimal contribution from the third population in this case.

#### S.6 Comparison of MDF and JR connectivity parameter spaces

By setting the parameters to be the same between JR and MDF, we compare the connectivity parameter space of the two models (Fig. S9).

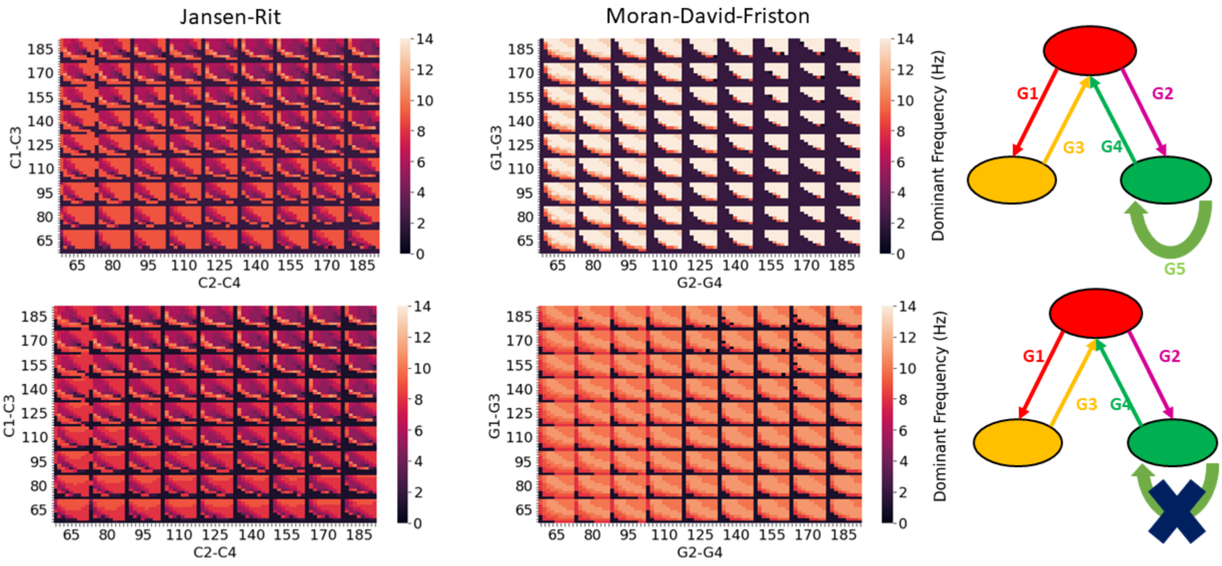


**Figure S9.** ***Connection strength parameter spaces for JR and MDF with similar parameter settings.*** In the top, for MDF $\gamma_5 = 16$, and at the bottom $\gamma_5 = 0$. The general shape of the dynamics is very similar between the two, suggesting that the effects of connectivity are the same. However, MDF tends to generate oscillations of higher frequencies for identical connectivity parameter sets, even when the $\gamma_5$ connection is removed.

In the top row of Fig. S6, we compare JR against MDF with the self-inhibitory connection. We observe a similar triangular boundary shape within which the system oscillates. However, MDF tends oscillate at higher frequency that exceed the alpha range (Fig. S6, MDF top row, colors are brighter than JR). When the self-inhibitory connection is removed in MDF (Fig. S6, MDF bottom row), the system now oscillates at the alpha frequency. It does not present lower frequencies, such as those in the JR model where we have slower oscillations. Thus, MDF seems to oscillate at higher frequencies than JR. Nonetheless, we observe that the two models share this similar triangular shape with non-oscillatory behavior when $C_3$ and $C_4$ are too low, suggesting similar global dynamics. The main conclusion drawn from this analysis is that the self-inhibitory connection introduced in MDF grants the model the ability to generate oscillations at a higher frequency than alpha, a more challenging capability compared to JR.

#### S.7 4D JR connectivity analysis

In the JR model, our focus was specifically on $C_3$ ($P \rightarrow I$) and $C_4$ ($I \rightarrow P$) as the E-I loop, but there is also the interaction between excitatory interneurons and pyramidal cells ($C_1$ ($P \rightarrow E$) and $C_2$ ($E \rightarrow P$)) to consider. Typically, the ratio between these values is varied. By simulating time series for different values of $C$ with the standard ratio values ($C_1 = C$, $C_2 = 0.8 * C$, $C_3 = 0.25 * C$ and $C_4 = 0.25 * C$), we can infer that increasing values of C lead to a decrease in the frequency of oscillation up to a certain point (Fig. S5), which concurs with results from Jansen and Rit (1995).

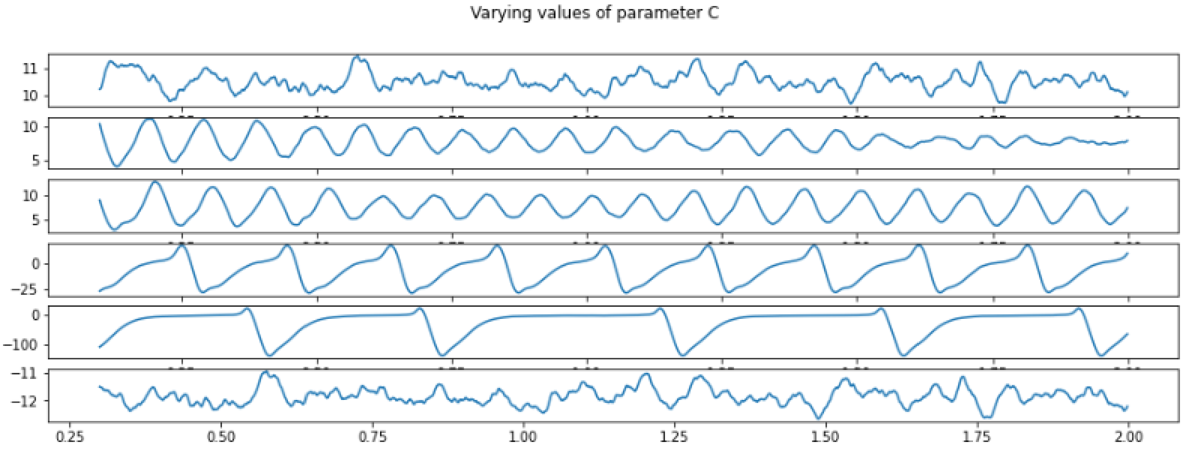


**Figure S5.** ***Simulated time series of JR for different connectivity C values.*** From top to bottom: $C = 68$, $C = 128$, $C = 135$, $C = 270$, $C = 675$, $C = 1350$. As connectivity values increase, the frequency of oscillations decreases up to $C = 675$.

Changes in the frequency of oscillation as a function of connectivity ratios are presented in the form of 4D heatmaps in a 2D space (Fig. S6). The general trend observed is that higher connectivity values result in slower oscillations, as expected.

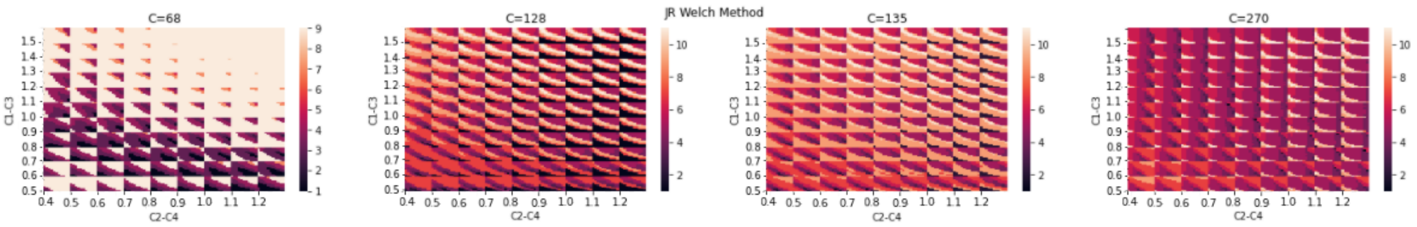


**Figure S6.** ***Variation in the frequency of oscillation as a function of connection strength for JR for different C values.*** From left to right: $C = 68$, $C = 128$, $C = 135$ and $C = 270$. The outer axes $C_1 - C_2$ represent the excitatory loop, while the inner axes $C_3 - C_4$ represent the inhibitory loop. The results correlate with what is observed in the time series. The parameter space is obtained by changing the ratio of each connection (i.e: $C1 = 0.8$ corresponds to $C1 = 0.8 * C$)

To observe the different trends that emerge, we focused on the case where $C$ equals 135 and investigated the different possible combinations of parameters on the outer and inner axes (Fig. S7).

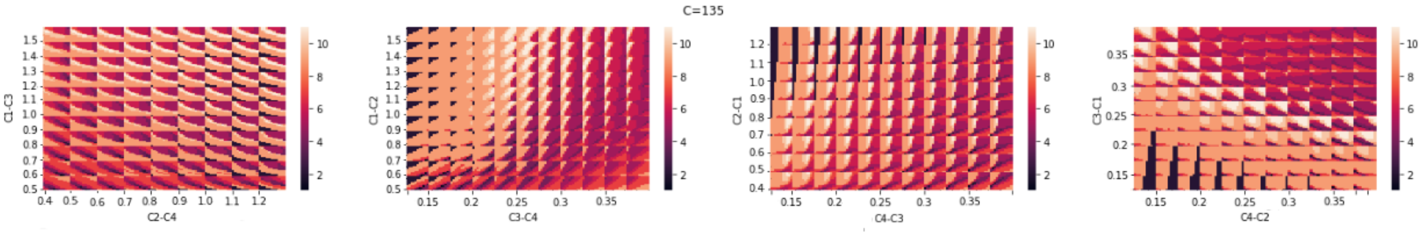


**Figure S7.** ***Connection strength parameter spaces for Jansen-Rit in different combinations with C=135.*** Each combination reveals a distinct pattern, aiding in visualizing the relationships among all the connectivity parameters. From left to tight: 1) Outer axes $C_1 - C_2$ Inner axes $C_3 - C_4$; 2) Outer axes $C_1 - C_3$; Inner axes $C_2 - C_4$; 3) Outer axes $C_2 - C_3$; Inner axes $C_1 - C_3$; 4) Outer axes $C_3 - C_4$; Inner axes $C_1 - C_2$

Clear patterns emerge in two different cases. When $C_3$ ($P \rightarrow I$) and $C_4$ ($I \rightarrow P$) are on the outer axes, a continuous change in the frequency of oscillation is observed. Similarly, when comparing $C_1$ ($P \rightarrow E$) against $C_3$ ($P \rightarrow I$), a concrete pattern is evident, with more pronounced changes in the frequency of oscillation when $C_3$ is altered. These results reinforce the idea that the main loop influencing the frequency of oscillation is the interaction between the pyramidal and inhibitory populations, raising the question of whether adding an additional excitatory population is truly necessary, even though it would be more biologically realistic.

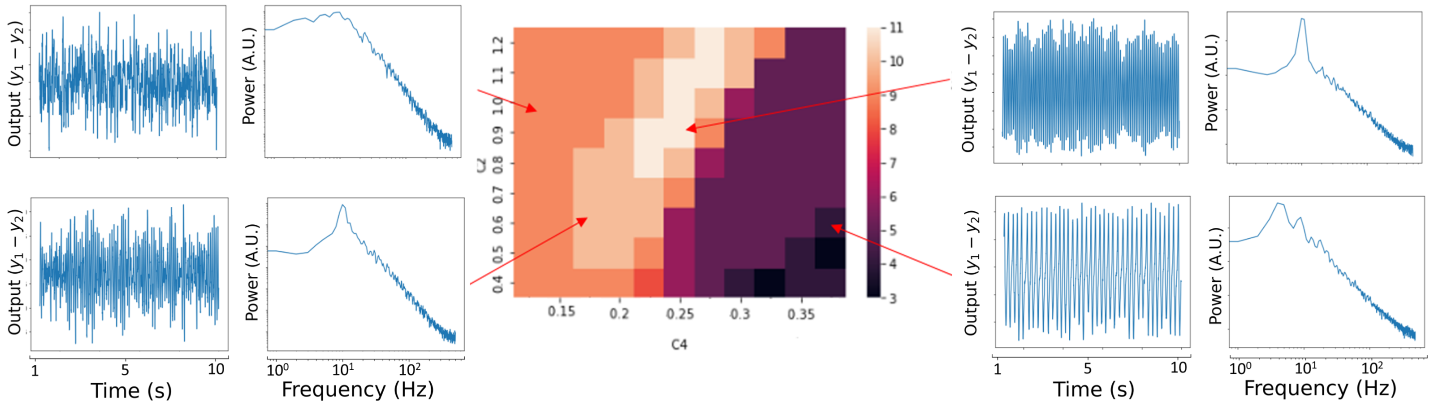


**Figure S8.** ***Connection strength parameter space for*** $C_2$ ***(***$E \rightarrow P$***) and*** $C_4$ ***(***$I \rightarrow P$***) in JR.*** Higher values of $C_4$ lead to a decrease in rhythmic oscillations. The highest frequency of oscillation occurs when $C_4$ is at a ratio of 0.25, and $C_2$ is around 1.0. If $C_4$ is too low, a very noisy signal is generated.

#### S.8 3D parameter space with MDF

We simplified the 5-dimensional connection parameter space into a 3-dimensional representation for the MDF model, using its linearized version. Stability is assessed by looking at the system's poles within the transfer function of the system. The aim was to establish a parallel with the 3D 'xyz' corticocortical/corticothalamic/intrathalamic lumped gains reduced parameter space discussed in a number of studies using the RRW model (although for reasons of space we have not focused on that aspect of RRW in the present paper Robinson et al., 2002, 2005; Roberts and Robinson, 2012; Breakspear et al., 2006; Abeysuriya et al., 2015), and determine the effects of the loops on the dynamics of the MDF model.

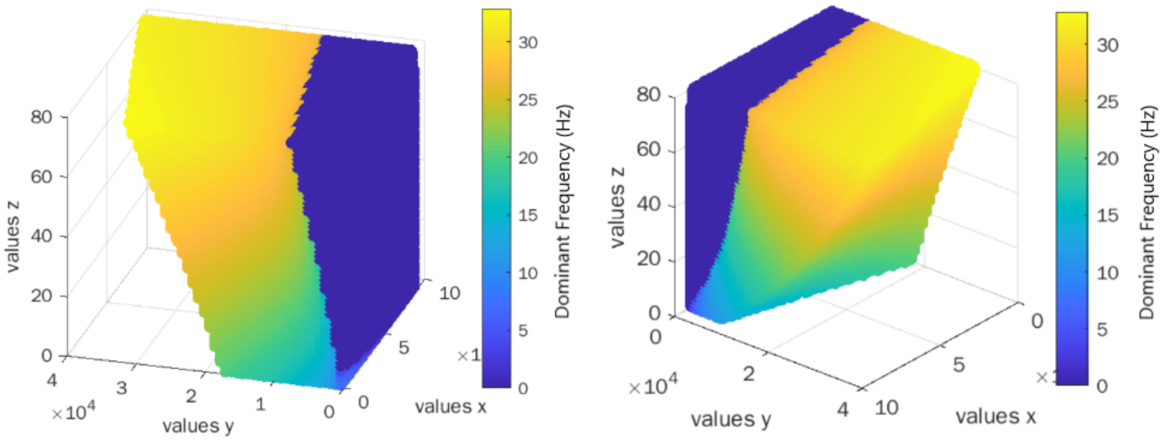


**Figure S10.** ***Visualization of dynamical regimes of the MDF model in a 3D setting using the linearized expression.*** The x-axis corresponds to effect of the excitatory loop ($\gamma_1 * \gamma_2$); the y-axis represents the effect of the inhibitory loop ($\gamma_3 * \gamma_4$); and the z-axis is the effect of the self-inhibitory loop ($\gamma_5$). As $\gamma_5$ values increase, the system tends to oscillate at a higher frequency.

The aim here is to easily visualize the regions of stability and dynamics as a function of the 'loops', rather than a single connectivity parameter. As expected, with the increase in the self-inhibitory connection (z-axis), the dominant frequency of oscillation gradually shifts from theta to alpha and then to the beta range.

#### S.9 Full Model Equations

Partial and diagrammatic presentations of the differential equations for each of the four models are given in Figs. 4-7. In this final Supplementary section, we provide the complete differential equations for each model, as well as tables describing the model parameters and state variables.

##### Jansen-Rit model equations

The differential equations for the JR model are

$$\dot{y}_0(t) = y_3(t) \tag{29}$$

$$\dot{y}_3(t) = AaS[y_1(t) - y_2(t)] - 2ay_3(t) - a^2y_0(t) \tag{30}$$

$$\dot{y}_1(t) = y_4(t) \tag{31}$$

$$\dot{y}_4(t) = Aa(p(t) + C_2S[C_1y_0(t)]) - 2ay_4(t) - a^2y_1(t) \tag{32}$$

$$\dot{y}_2(t) = y_5(t) \tag{33}$$

$$\dot{y}_5(t) = BbC_4S[C_3y_0] - 2by_5(t) - b^2y_2(t) \tag{34}$$

Here and in the rest of this paper we have maintained the same notation as in Jansen and Rit (1995) where $y_0$, $y_1$, and $y_2$ correspond to the outputs of the pyramidal, excitatory, and inhibitory PSP block, respectively. $p(t)$ represents the external input applied to the system, usually noise. $A$ and $B$ define the maximum amplitude of excitatory and inhibitory PSP, respectively. $a$ and $b$ represent the collective effect of the inverse of the time constant of the passive membrane and the entirety of the spatially dispersed delays within the dendritic network for the excitatory and inhibitory populations, respectively. $C_1$ to $C_4$ are the connectivity constants.

For the connectivity parameters, we wanted to mention that $C_1$ and $C_3$ slightly differ from $C_2$ and $C_4$ in the mathematical expression. The JR model assumes equal synaptic input from the pyramidal cell population to the other two populations, setting these constants to 1. In contrast, the synaptic coefficients at the excitatory and inhibitory dendrites are varied (corresponding to $C_1$ ($P \to E$) and $C_3$ ($P \to I$)). Conversely, for pyramidal cells, the synaptic coefficients at their dendrites remain fixed (1 and -1 for excitatory and inhibitory interneurons, respectively), and excitatory and inhibitory neurons synapse onto pyramidal cells differently (represented by $C_2$ ($E \to P$) and $C_4$ ($I \to P$). Therefore, $C_1$ and $C_3$ function as synaptic coefficients, while $C_2$ and $C_4$ serve as connectivity constants, as illustrated in the detailed schematic. Mathematically, this means that $C_1$ and $C_3$ are applied within the nonlinear function, while $C_2$ and $C_4$ are applied outside. However, in practical terms, all these parameters are described as connectivity parameters and can be considered analogous and interrelated. Furthermore, all the values are scaled by a global connectivity parameter. See Cook et al. (2021) for a further explanation of this nuanced aspect of the JR model system.

| Symbol | Description | Value |
|---|---|---|
| $e_0$ | Firing rate at threshold | 2.5 $s^{(-1)}$ |
| $V_0$ | Firing threshold | 6 mV |
| r | Slope reflecting the variance of firing thresholds within the population | 0.56 $mV^{(-1)}$ |
| A | Maximum amplitude of excitatory PSP (EPSP) | 3.25 mV |
| B | Maximum amplitude of inhibitory PSP (IPSP) | 22 mV |
| a and b | Lumped representation of the sum of the reciprocal of the time constant of passive membrane and all other spatially distributed delays in the dendritic network | a = 100 $s^{(-1)}$<br>b = 50 $s^{(-1)}$ |
| $C_1$ | Connectivity constant: Represents the number of synapses made by the feed forward neurons to the dendrites of the excitatory feedback loop | $C = C_1$<br>135 |
| $C_2$ | Connectivity constant: Proportional to the number of synapses made by the excitatory feedback loop to the dendrites of the feedforward neurons | $C_2 = 0.8C$ |
| $C_3$ | Connectivity constant: number of synapses made by the feedforward neurons to the dendrites of the inhibitory feedback loop | $C_3 = 0.25C$ |
| $C_4$ | Connectivity constant: Proportional to the number of synapses made by the inhibitory feedback loop to the dendrites of the feedforward neurons | $C_4 = 0.25C$ |
| P(t) | External pulse density consisting of activity originating from adjacent and more distant cortical columns and from subcortical structures (e.g. thalamus) | Uniform noise (or normal, constant) |

**Table 4.** ***JR parameters with biological descriptions and corresponding values to generate alpha rhythm***

#### Moran-David-Friston model equations

The form of the differential equations for the MDF model are

$$\dot{\nu}_1 = i_1 \quad (35)$$

$$\dot{i}_1 = \kappa_e H_e(\gamma_1 S(\nu_6 - a) + u) - 2\kappa_e i_1 - \kappa_e^2 \nu_1 \quad (36)$$

$$\dot{\nu}_2 = i_2 \quad (37)$$

$$\dot{i}_2 = \kappa_e H_e \gamma_2 S(\nu_1) - 2\kappa_e i_2 - \kappa_e^2 \nu_2 \quad (38)$$

$$\dot{\nu}_3 = i_3 \quad (39)$$

$$\dot{i}_3 = \kappa_i H_i \gamma_4 S(\nu_7) - 2\kappa_i i_3 - \kappa_i^2 \nu_3 \quad (40)$$

$$\dot{\nu}_6 = i_2 - i_3 \tag{41}$$
$$\dot{\nu}_4 = i_4 \tag{42}$$
$$\dot{i}_4 = \kappa_e H_e \gamma_3 S(\nu_6) - 2\kappa_e i_4 - \kappa_e^2 \nu_4 \tag{43}$$
$$\dot{\nu}_5 = i_5 \tag{44}$$
$$\dot{i}_5 = \kappa_i H_i \gamma_5 S(\nu_7) - 2\kappa_i i_5 - \kappa_i^2 \nu_5 \tag{45}$$
$$\dot{\nu}_7 = i_4 - i_5 \tag{46}$$

The $v_i$ values represent the membrane potential of the subpopulations and $i_i$ denoting their current. Specifically, $v_1$ and $i_1$ describe the excitatory interneurons, $v_{2,3,6}$ and $i_{2,3}$ the pyramidal cells, and finally $v_{4,5,7}$ and $i_{4,5}$ the inhibitory interneurons. The $\gamma_i$ values are the connection strengths between the populations. $H_e$ and $\kappa_e$ are the maximum amplitude and the rate constant associated with EPSP, respectively. Similarly, $H_i$ and $\kappa_i$ represent the same parameters for the IPSP.

| Symbol | Description | Value |
|---|---|---|
| $\rho_1$ | For shape of sigmoid: Can straighten more or less the slope | 2 |
| $\rho_2$ | For position of sigmoid: Can shift the curve right or left | 1 |
| $H_e$ | Maximum amplitude of excitatory PSP (EPSP) | 10 mV |
| $H_i$ | Maximum amplitude of inhibitory PSP (IPSP) | 22 mV |
| $\kappa_e$ and $\kappa_i$ | Lumped representation of the sum of the rate constants of passive membrane and other spatially distributed delays in the dendritic tree | $\kappa_e = 250$ s$^{(-1)}$ $\kappa_i = 62.5$ s$^{(-1)}$ |
| $\gamma_1$ | Coupling strength: Between pyramidal cells and macrocolumn u (in excitatory spiny cells in granular layer) | 128 |
| $\gamma_2$ | Coupling strength: Between excitatory spiny cells in granular layer and pyramidal cells | 128 |
| $\gamma_3$ | Coupling strength: Between pyramidal cells (excitatory) and inhibitory interneurons | 64 |
| $\gamma_4$ | Coupling strength: Between inhibitory interneurons and pyramidal cells | 64 |
| $\gamma_5$ | Coupling strength: Inhibitory-Inhibitory coupling (recurrent connection) | 1 |

**Table 5.** ***MDF parameters with biological descriptions and corresponding values to generate alpha rhythm***

#### Liley-Wright model equations

For the LW model, the differential equations are

$$\begin{aligned}
\tau_e \dot{V}_e(t) &= V_e^{rest} - V_e(t) + \psi_{ee}(V_e(t))I_{ee}(t) + \psi_{ie}(V_e(t))I_{ie}(t) && (47)\\
\tau_i \dot{V}_i(t) &= V_i^{rest} - V_i(t) + \psi_{ei}(V_i(t))I_{ei}(t) + \psi_{ii}(V_i(t))I_{ii}(t) && (48)\\
\dot{I}_{ee} &= U_{ee} && (49)\\
\dot{U}_{ee} &= -2\gamma_e U_{ee}(t) - \gamma_e^2 I_{ee}(t) + \Gamma_e \gamma_e e(N_{ee}^{\beta} S(V_e(t)) + p_{ee}(t)) && (50)\\
\dot{I}_{ei} &= U_{ei} && (51)\\
\dot{U}_{ei} &= -2\gamma_e U_{ei}(t) - \gamma_e^2 I_{ei}(t) + \Gamma_e \gamma_e e(N_{ei}^{\beta} S(V_e(t)) + p_{ei}(t)) && (52)\\
\dot{I}_{ie} &= U_{ie} && (53)\\
\dot{U}_{ie} &= -2\gamma_i U_{ie}(t) - \gamma_i^2 I_{ie}(t) + \Gamma_i \gamma_i e(N_{ie}^{\beta} S(V_i(t))) && (54)\\
\dot{I}_{ii} &= U_{ii} && (55)\\
\dot{U}_{ii} &= -2\gamma_i U_{ii}(t) - \gamma_i^2 I_{ii}(t) + \Gamma_i \gamma_i e(N_{ii}^{\beta} S(V_i(t))) && (56)
\end{aligned}$$

$N_{xx}$ are the inter- and intra-connectivities between the two populations. $p_{ei}$ and $p_{ee}$ are the external inputs. $I_{xx}$ are the postsynaptic potentials, and $V_{xx}$ are the soma membrane potentials. $\Gamma_{e,i}$ and $\gamma_{e,i}$ are the peak amplitude and rate constant PSPs for excitatory and inhibitory population, respectively. The model also includes passive membrane time constants represented by $\tau_{e,i}$, mean resting membrane potentials $V_{e,i}^{r}$, and mean equilibrium potentials $V_{e,i}^{eq}$.

| Symbol | Description | Value |
|---|---|---|
| $S_{(e,i)}^{max}$ | Excitatory/Inhibitory population mean maximal firing rates | 500, 500 $s^{(-1)}$ |
| $\mu_{(e,i)}$ | Excitatory/Inhibitory population thresholds (spike threshold) | -50, -50 mV |
| $\sigma_{(e,i)}$ | Standard deviation for spike-threshold in excitatory/inhibitory population | 5, 5 mV |
| $\Gamma_e$ | Excitatory postsynaptic potential peak amplitude (at the site of synaptic activation) | 0.71 mV |
| $\Gamma_i$ | Inhibitory postsynaptic potential peak amplitude (at the siyte of synaptic activation) | 0.71 mV |
| $\gamma_{(e,i)}$ | Excitatory/Inhibitory postsynaptic potential rate constant | 300, 65 $s^{(-1)}$ |
| $\tau_{(e,i)}$ | Passive membrane decay time constant | 0.094, 0.042 s |
| $V_{(e,i)}^{r}$ | Mean resting membrane potential | -70, -70 mV |
| $V_{(e,i)}^{eq}$ | Mean equilibrium potential associated with excitation or inhibition | 45, -90 mV |
| $N_{(ee,ei)}^{\beta}$ | Total number of connections that a cell of type e, i receives from excitatory cells via intra-cortical fibres (Weight connections) | 3000, 3000 |
| $N_{(ie,ii)}^{\beta}$ | Total number of connections that a cell of type e,i receives from inhibitory cells via intra-cortical connections (Weight connections) | 500, 500 |
| $p_{(ee,ei)}$ | Excitatory extra-cortical input to excitatory, inhibitory cells | 3.46, 5.07$s^{(-1)}$ |
| $p_{(ie,ii)}$ | Inhibitory extra-cortical input to excitatory, inhibitory cells | 0, 0 $s^{(-1)}$ |

**Table 7.** ***LW parameters with biological descriptions and values to generate alpha rhythm***

#### Robinson-Rennie-Wright model equations

Finally, the differential equations of the RRW are as follows

$$
\begin{aligned}
\frac{dV_e}{dt} &= \dot{V}_e && (57)\\
\frac{d\dot{V}_e}{dt} &= \alpha\beta[\nu_{ee}\phi_e + \nu_{ei}S(V_e) + \nu_{es}S(V_s(t - t_0/2)) - (\frac{1}{\alpha} + \frac{1}{\beta})\dot{V}_e - V_e] && (58)\\
\frac{dV_s}{dt} &= \dot{V}_s && (59)\\
\frac{d\dot{V}_s}{dt} &= \alpha\beta[\nu_{se}\phi_e(t - t_0/2) + \nu_{sr}S_r(V_r) + \nu_{sn}\phi_n - (\frac{1}{\alpha} + \frac{1}{\beta})\dot{V}_s - V_s] && (60)\\
\frac{dV_r}{dt} &= \dot{V}_r && (61)\\
\frac{d\dot{V}_r}{dt} &= \alpha\beta[\nu_{re}\phi_e(t - t_0/2) + \nu_{rs}S(V_s) - (\frac{1}{\alpha} + \frac{1}{\beta})\dot{V}_e - V_r] && (62)\\
\frac{d\phi_e}{dt} &= \dot{\phi}_e && (63)\\
\frac{d\dot{\phi}_e}{dt} &= \gamma_e^2[S(V_e) - \frac{2}{\gamma_e}\dot{\phi}_e - \phi_e] && (64)
\end{aligned}
$$

with $V_e$, $V_r$, and $V_s$ representing the potential of the cortical population, the reticular nucleus, and the relay nuclei, respectively. $\nu_{xx}$ denote the connection strengths parameters. $\alpha$ and $\beta$ refer to the decay and rise time of the impulse response, representing the dendritic rate. $t_0$ is the conduction delay between thalamic and cortical projections. Finally, $\gamma_e$ stands for the cortical damping rate, which is exclusively applied to the cortical population. This final differential equation for determining $\phi_e$ is related to the PDE damped wave equation, used to consider spatial variations (Robinson et al., 1997). However, in the case of spatial uniformity, the wave equation simplifies to an ODE (Zhao and Robinson, 2015).

| Symbol | Description | Value |
|---|---|---|
| $Q_{max}$ | Maximum attainable firing rate of individual neurons | 340 s$^{(-1)}$ |
| $\sigma'\pi\sqrt{3}$ | Standard deviation of the threshold distribution in the neural population | $3.8{*}\pi\sqrt{3} \approx$ 5.9 mV |
| $\theta$ | Mean firing threshold | 12.92 mV |
| $\gamma_e$ | Cortical damping rate (Axonal velocity/Range) | 116 $s^{(-1)}$ |
| $1/\alpha$ | Decay time (of impulse response, dendritic rate) | 83.33 $s^{-1}$ |
| $1/\beta$ | Rise time (of impulse response, dendritic rate) | 769.23 $s^{-1}$ |
| $t_0$ | Corticothalamic loop delay (Loop distance/Axonal velocity = conduction delay through thalamic nuclei and projections) | 80 ms |
| $v_{ee}$ | $N_{ee}s_{ee}$: Mean number of synapses X strength of the response to a unit signal | 3.03 mVs |
| $-v_{ei}$ | $-N_{ei}s_{ei}$ | 6.00 mVs |
| $v_{es}$ | $N_{es}s_{es}$ | 2.06 mVs |
| $v_{se}$ | $N_{se}s_{se}$ | 2.18 mVs |
| $-v_{sr}$ | $-N_{sr}s_{sr}$ | 0.83 mVs |
| $v_{re}$ | $N_{re}s_{re}$ | 0.33 mVs |
| $v_{rs}$ | $N_{rs}s_{rs}$ | 0.03 mVs |
| $v_{sn}$ | $N_{sn}s_{sn}$ | 0.98 mVs |

**Table 6.** ***RRW parameters with biological descriptions and values to generate alpha rhythm***

#### S.10 Additional background literature

For interested readers, we provide here some further background literature on early history of NPMs and other alpha theories:

##### Tracing the roots of NPMs: early history

The notion of neural *masses* was introduced in various forms during the 1950s and 1960s (Beurle, 1956; Griffith, 1963), and consolidated in the 1970s primarily through the highly influential work of Freeman, Wilson & Cowan, Amari, and Nunez. It was Freeman who originally used the term 'neural mass action model' (Freeman, 1972a,b, 1975), articulating many of the neurobiological and mathematical fundamentals as they are understood today in a wide-reaching monograph on the subject (Freeman, 1975). Here, Freeman also develops the theory of 'K-sets' which are based on a hierarchy of interacting sets of neural populations or masses, and used to model neural population dynamics with ordinary differential equations (ODEs) to simulate mesoscopic local field potentials (Deschle et al., 2021). The levels are designated as K0, KI, KII, and KIII, with the K0 set corresponding to a model characterized by non-interactive collections of neurons with globally common inputs and outputs, KI to pairs of interacting K0 sets, and so on. Freeman's research on the olfactory bulb and prepyriform cortex of cats and rabbits (Freeman, 1979, 1975) provides valuable experimental data that has been used to define mathematical formulations and parameter settings in many NMMs, which is further discussed in section 3.2.4. Furthermore, Freeman's contributions on the use of the sigmoidal operator for mapping membrane potential to firing rate remains a critical component of many NMMs, the validity of which will be elaborated on in section 4.2. Even though Freeman coined the term neural masses and laid much of the groundwork, many of the core mathematical principles of NMMs were first proposed in the work of Wilson & Cowan (WC; Wilson and Cowan, 1972), which itself builds upon earlier work by Beurle (1956). WC's implementation introduced and solidified an approach to modelling neural dynamics and brain function. This approach consists of analyzing the collective properties of a large number of neurons using methods from statistical mechanics rooted in the mean-field framework (Destexhe and Sejnowski, 2009; Chow and Karimipanah, 2020). By omitting potential spatial arrangement of synaptic connections, their model offers a minimalistic NMM representation that has been leveraged to develop several simple yet biophysically plausible models (eg Kilpatrick, 2013; Sanz-Leon et al., 2015). As shown in Fig. S11, the canonical WC model consists of two neural masses with one excitatory and one inhibitory population (Wilson and Cowan, 1972; Sanz-Leon et al., 2015). Two nonlinear ODEs describe the dynamics of those two synaptically coupled populations in the neocortex (Nakagawa et al., 2014; Cowan et al., 2016). The WC system is thus a coarse-grained description of the overall activity and mesoscale neuronal network structure of a patch of (usually cortical) tissue, as is typical of NPMs. By varying the connectivity strength and the input strength to each population, it is possible to generate a diversity of dynamical behaviors that are characteristic of observed activity in the brain, such as multistability, oscillations, traveling waves, and spatial patterns (Kilpatrick, 2013).

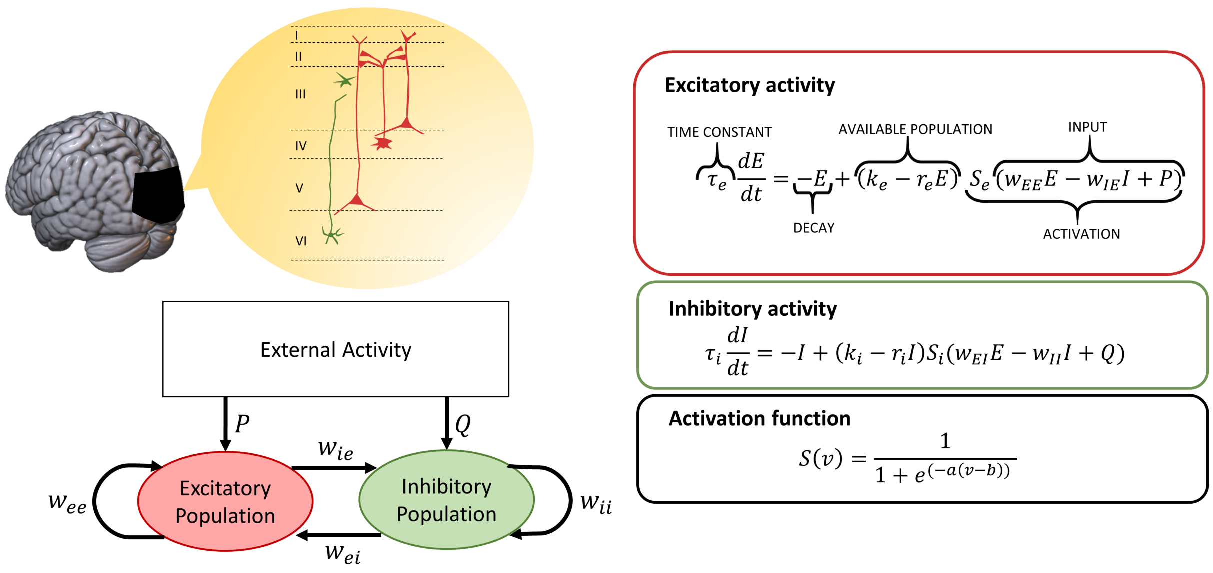


**Figure S11.** ***Wilson-Cowan model topography and mathematical expression.*** The model aims to represent a cortical column within the brain, consisting of an excitatory and an inhibitory population. These two connected populations each have a self-connection and external activity as input. Dynamics are expressed with nonlinear ordinary differential equations which are shown on the right for each neural population. Nonlinearity is introduced with the sigmoidal operator corresponding to the activation function.

A simplified version of the WC equations shown in Fig. 3 has been previously implemented by Abeysuriya et al. (2018) in a network of neural masses to generate alpha oscillations. These two populations are described as follows:

$$\tau_e \frac{dE(t)}{dt} = -E(t) + S(w_{ee}E(t) + w_{ie}I(t) + P + \epsilon(t)) \quad (65)$$

$$\tau_i \frac{dI(t)}{dt} = -I(t) + S(w_{ei}E(t) + \epsilon(t)) \quad (66)$$

where $E$ and $I$ represent the activity of the excitatory and inhibitory neural populations in the form of mean firing rates, $\tau_{e/i}$ are the excitatory/inhibitory time constants, $w_{ab}$ are the local connection strengths from population $a$ to population $b$, $P$ is a constant external input to the excitatory neural population, and $\epsilon$ is a noise signal added to the system. The studied NMMs share similar parameters, with some variations such as the use of membrane potential instead of firing rates as the state variable, and the concatenation of the external input and noise term into a single variable.

Concurrently to WC and Freeman, Lopes da Silva and colleagues developed a point-process model of EEG alpha rhythm generated with a corticothalamic loop (Lopes da Silva et al., 1974). Specifically, these authors proposed a negative feedback loop between excitatory thalamocortical relay cells and inhibitory thalamic reticular neurons as the basis for generating certain brain rhythms, in a manner similar to the interacting E and I populations in the WC model. By applying linear systems analysis to investigate the influence of physiological parameters on neural periodic patterns, they established a novel approach to studying oscillatory dynamics in theoretical neuroscience that relied on analytical power spectra. The Lopes da Silva model had a substantial impact on subsequent corticothalamic models and linear analysis tools (Cona et al., 2014; Bhattacharya et al., 2011).

A few years later, Zetterberg et al. (1978) built an extension of the model by adding a second cortical excitatory population in order to separately account for pyramidal cells and excitatory interneurons. Their work was then reprised and further popularized by Jansen and Rit (1995). In the JR model, each neural population is described in two steps: a transformation of the incoming average pulse density of action potentials into an average postsynaptic membrane potential, followed by a sigmoidal function to perform the inverse conversion. Over the years, several extended versions of JR have been proposed (Wendling et al., 2000; David and Friston, 2003; Zavaglia et al., 2006; Sotero et al., 2007), - including Moran et al., where they focused on steady-state spectral responses with a linearized approximation of the model (Moran et al., 2007). Contemporaneous with these early conceptualizations and formulations of NMMs in the 1970s was the introduction of NFMs by Amari, Wilson & Cowan, Nunez, and others. The 'brain wave equation' model of (Nunez, 1974) is particularly important here as it was the first to attempt to describe neural activity across the entire cerebral cortex with an evolution in both time and space. This work was a major influence for several macroscale NFM formulations in the 1990s (Jirsa and Haken, 1996; Wright and Liley, 1996; Robinson et al., 1997). The latter of these which was then extended in 2001 to include the thalamus, and subsequently used to investigate a wide range of brain states including sleep (Robinson et al., 2005; Abeysuriya et al., 2014), epileptic seizures (Zhao and Robinson, 2015; Breakspear et al., 2006), evoked responses (Kerr et al., 2008), functional connectivity (Robinson, 2014), and alpha rhythms (Robinson et al., 2002, 2005).

For a more detailed timeline and review on the development of NPMs and whole brain modelling in general, we refer the reader to Griffiths et al. (2022) and Chow and Karimipanah (2020). The early mathematical models reviewed there and above laid the groundwork for most NPM formulations used in theoretical neuroscience today. In particular, they form the basis for the four most widely studied models of the EEG alpha rhythm - Jansen-Rit (JR), Moran-David-Friston (MDF), Liley-Wright (LW) and Robinson-Rennie-Wright (RRW).

#### Pacemaker vs. Local Network vs. Global Network Alpha theories

As indicated in the main text, the theories of alpha in the literature can be grouped under three categories: *pacemaker*, *local network*, and *global network* theories. For reasons of space we discuss only the second of these in this article. The following provide some further notes and references for the interested reader on the other two. The pacemaker theory suggests that intrinsic alpha oscillations are generated either in the thalamus, driven by pulvinar or and/or the lateral geniculate nucleus (Saalmann et al., 2012; Lőrincz et al., 2009; Hughes et al., 2011) or in the cortex, originating from the pyramidal cells located in layer V (Lopes da Silva, 1991; Connors and Amitai, 1997; Bollimunta et al., 2008). However, pacemaker theories in general suffer from several severe limitations (see Nunez et al. 2006 for an extensive discussion of this). For instance, pacemaker cells such as putative thalamic nuclei, if they exist, would have to function in a relatively autonomous fashion, having a highly restricted input from other oscillatory brain regions - a notion that has been critically questioned on anatomical grounds (Lopes da Silva, 1998; Steriade, 2005). Additionally, there are certain global EEG phenomena that remain unexplained, including the relative frequencies of major rhythms and sleep-wave variations. The second category, 'local network' theories, propose that alpha rhythms are produced by interactions between excitatory and inhibitory neural populations with dendritic response functions and saturating nonlinearities (Valdés-Hernández et al., 2010). Finally, 'global network' theories posit that alpha rhythms are generated by large-scale networks rather than local circuits within a localized brain region. By disregarding complex dendritic response functions and finite intracortical propagation, models with a primary emphasis on global dynamics rely heavily on the propagation delays between distant anatomical structures to shape their dynamics (Nunez and Cutillo, 1995; Nunez and Srinivasan, 2006; Valdés-Hernández et al., 2010).
